# Intraspecific difference among herbivore lineages and their host-plant specialization drive the strength of trophic cascades

**DOI:** 10.1101/722140

**Authors:** Arnaud Sentis, Raphaël Bertram, Nathalie Dardenne, Jean-Christophe Simon, Alexandra Magro, Benoit Pujol, Etienne Danchin, Jean-Louis Hemptinne

## Abstract

Trophic cascades—the indirect effect of predators on non-adjacent lower trophic levels—are important drivers of the structure and dynamics of ecological communities. However, the influence of intraspecific trait variation on the strength of trophic cascade remains largely unexplored, which limits our understanding of the mechanisms underlying ecological networks. Here we experimentally investigated how intraspecific difference among herbivore lineages specialized on different host plants influences the strength of trophic cascade in a terrestrial tritrophic system. We found that the occurrence and strength of the trophic cascade are strongly influenced by herbivores’ lineage and host-plant specialization but are not associated with density-dependent effects mediated by the growth rate of herbivore populations. Our findings stress the importance of intraspecific heterogeneities and evolutionary specialization as drivers of the strength of trophic cascades and underline that intraspecific variation should not be overlooked to decipher the joint influence of evolutionary and ecological factors on the functioning of multi-trophic interactions.

## Introduction

Predators strongly influence the structure and function of ecological communities by influencing prey density, distribution, and behavior which, in turn, have cascading effects on lower trophic levels (Sih et al. 1985; Beckerman, Uriarte & Schmitz 1997; Shurin et al. 2002; Schmitz, Krivan & Ovadia 2004; Suraci et al. 2016). This indirect effect of predators on non-adjacent lower trophic levels, the so-called “trophic cascades,” are frequently observed in both aquatic and terrestrial ecosystems (Halaj & Wise 2001; Borer et al. 2005; Bruno & O’Connor 2005; Wu et al. 2011; Sanders, Kehoe & van Veen 2015). Trophic cascades are important drivers of the structure and dynamics of populations, communities, and ecosystems (Ripple et al. 2016) and have several implications for theoretical ecology, conservation biology, and ecosystem management (Post et al. 1999; Hulot et al. 2000; Estes et al. 2011). For instance, Delvin et al. (2015) showed that stocking fishes in fishless lakes decreases by a factor 10 the efflux rates of methane—an important greenhouse gas—by reducing zooplankton abundance, which in turn increases the abundance of methanotrophic bacteria. In another study, Schmitz et al. (2017) showed that the composition of the arthropod predator community and associated cascading effects on plant communities explain 41% of the variation in soil carbon retention across a human land-use gradient. Given the importance of trophic cascades, a major issue in ecology and conservation is to determine when and where trophic cascades occur, and what are the factors and mechanisms underpinning their strength.

Although the existence of trophic cascades has been widely demonstrated (Schmitz 2003; Romero & Koricheva 2011), most studies focused on whether cascades are more likely or stronger in some systems than others (Pace et al. 1999; Shurin et al. 2002), leading to a wealth of predictions about the relative strength of predator effects on plants among ecosystems with a particular focus on aquatic versus terrestrial ecosystems (Shurin et al. 2002; Borer et al. 2005). Although comparing the strength of trophic cascade among systems is valuable, little is known about what causes variation in the magnitude of cascading effects within or among systems. Most prior studies assumed that species identity or mean trait values adequately represent species interactions and their effects on community dynamics. This assumption is puzzling because it ignores the considerable intraspecific variation of traits (Benesh & Kalbe 2016), thereby overlooking a potentially important determinant of species interactions, community structure and dynamics, as well as evolutionary responses to selective pressures (Roff 1997; Bolnick et al. 2011; Violle et al. 2012). Thus, knowledge of how intraspecific differences in phenotypic traits modulates bi- and tri-trophic interactions is crucial for better understanding and predicting the occurrence and strength of trophic cascades.

Different hypotheses have been proposed to explain variations in the occurrence and strength of trophic cascades in various types of ecosystems (Hulot et al. 2000; Polis et al. 2000; Borer et al. 2005). Hypotheses linked to the spatial heterogeneity of habitats, food web linearity or system productivity have received little support (Borer et al. 2005). On the other hand, the role of intraspecific trait variation remains largely unexplored. Recent studies showed that intraspecific variation in predator traits and behavior can influence the strength of trophic cascade (Clegg & Barlow 1982; Post et al. 2008; Jochum et al. 2012; Weis & Post 2013; Keiser et al. 2015; Royauté & Pruitt 2015; Start & Gilbert 2017). However, the role of intraspecific variation in herbivore traits and demography is still relatively unknown. It has been suggested that interspecific differences in plant anti-herbivore defenses (Schmitz et al. 2000; Mooney et al. 2010), predator hunting mode and consumer efficiency (e.g., low metabolic costs, high consumption rate and population growth rate) can significantly affect the strength of trophic cascade (Romero & Koricheva 2011). In particular, high predator or herbivore efficiency increases cascade strength via high consumption rate of herbivores by predators, and plants by herbivores (Strong 1992; Polis 1999; Borer et al. 2005). In other words, when consumers are efficient at consuming and converting their resources into new biomass, this translates in higher population growth rate that then strengthens the consumer impacts on the next trophic level. Therefore, the strength of trophic cascades should depend on the growth rate of herbivore populations in the absence of predators and on the predator efficiency at reducing herbivore density (Schmitz 1998; Borer et al. 2005). Density-mediated effects driven by intraspecific variation in herbivore population growth rate should thus affect trophic cascade strength: the faster the herbivore population growth, the stronger the trophic cascade. Consumer efficiency can vary among populations and food sources. Herbivore lineages can be specialized on different food sources and have evolved an ability to grow better on specific host plants. For instance, the pea aphid *Acyrthosiphon pisum* Harris (Homoptera: Aphididae) feeds on many Fabaceae species and forms host-plant-associated populations (“host races” or “biotypes”) that are genetically differentiated (Via 1999; Hawthorne & Via 2001; Peccoud et al. 2009) in a way that affects their population growth rate on different host plants (Via 1999; Via, Bouck & Skillman 2000; Hawthorne & Via 2001). We thus expected that host-plant specialization should lead to stronger trophic cascades when herbivores are adapted to their plant and efficient at converting them into new biomass.

In this study, we experimentally investigated the effects of herbivore intraspecific trait differences on trophic cascade strength using a broad bean–pea aphid–ladybeetle system. We conducted a full factorial laboratory experiment with six pea aphid clonal lineages (i.e. asexually reproducing aphid genetic lines) specialized either on alfalfa (*Alfalfa* biotype) or clover (*Clover* biotype) and exposed or not to a generalist predator. We first tested whether trophic cascade strength varied among aphid clonal lineages, and then investigated whether differences among lineages were best explained by density-mediated effects or by host-plant specialization. Our study highlights the importance of accounting for intraspecific differences and resource specialization to better understand and predict the strength of trophic cascades.

## Methods

### Biological system

The experimental system comprised a three levels food chain: the predatory ladybeetle *Harmonia axyridis* Pallas (Coleoptera: Coccinellidae), the pea aphid *A. pisum*, and the broad bean *Vicia faba* L. cv. Aquadulce. The broad bean *V. faba* is the universal legume host on which all pea aphid biotypes can feed and successfully develop (Peccoud et al. 2009). Approximately 200 ladybeetle adults *H. axyridis* were collected in October 2015 near Auzeville Tolosane (43°32’N, 1°29’E, South of France), brought to the laboratory, reared in 5000-cm3 plastic boxes, and fed three times a week an excess of pea aphids and pollen. Corrugated filter paper was added to each box to provide a suitable substrate for oviposition. *Harmonia axyridis* eggs were collected three times a week and neonate larvae were reared in 175-cm3 plastic boxes and fed pea aphids *ad libitum* until the beginning of the experiments. Stock colonies of 6 pea aphid clonal lineages (T9005, 10TV, T734, LL01, LSR1, and Oxford 683) were maintained in our laboratory at low density on broad bean grown from seeds (Ets Henrion s.a.; Belgium) in nylon cages (30 × 30 × 30 cm) for more than three months before the beginning of the experiments. All aphid lineages were free of any of the eight secondary symbionts reported in the pea aphid (Gauthier et al. 2015) (i.e. only harbour the obligate endosymbiont *Buchnera aphidicola*) to avoid potential confounding effects of variation in symbiont composition among aphid lineages. These lineages were selected from a large collection of clones maintained at INRAE Rennes and their symbiotic status was checked using diagnostic PCR as described in Peccoud et al. (2015). Three lineages (T9005, 10TV, and T734) were of the *Clover* biotype and three (LL01, LSR1, and Oxford 683) of the *Alfalfa* biotype. For each biotype, one of the tested lineages had a green color (T9005 and LL01) whereas the two other lineages were pink (10TV, T734, LSR1, Oxford 683). We used a standard set of seven microsatellite loci to confirm that each lineage represented a unique genotype (clone) and that each belonged to the aphid biotype corresponding to the plant from which it was collected (Peccoud et al. 2009). All insects and plants were maintained in air-conditioned chambers (Dagard^®^) at 21 ± 1°C, 50–60% relative humidity, and under a 16L:8D photoperiod. These experimental conditions ensure the pea aphid reproduces only by apomictic parthenogenesis (i.e., offspring are clones of their mother).

### Experimental design

In a full factorial laboratory experiment, we measured the effects of the 6 aphid clonal lineages and predators (presence or absence) on the fresh aboveground biomass and height of broad bean plants. At the onset of the experiment, three 8- days-old bean plants with two unfurled leaves were transplanted in 500 mL plastic pots containing 400 mL of fertilized soil substrate (^®^Jiffy substrates NFU 44-551), and then enclosed in transparent plastic cylinders (ø: 14 cm; h: 29 cm). They were watered every three days with 75 mL of tap water per pot. The top of the cylinder and two lateral openings were covered with mesh muslin for ventilation. Six parthenogenetic two-days-old adult female *A. pisum* were transferred to the upper leaves of the plants using a fine paintbrush and allowed to acclimatize and reproduce for 24 h. Then, one second instar *H. axyridis* larva was introduced into each experimental cylinder of the predation treatment. Ten days later, all aphids were collected using a fine paintbrush and counted under a stereoscopic microscope. The ladybeetle larvae were isolated in small Petri dishes (50 × 9 mm) and starved for 24h to empty their gut before being weighed with a micro-balance (10^−7^ g, SC2, Sartorius^®^). The plants were harvested, and their height and fresh aboveground biomasses measured. There were 20 replicates for each combination of aphid lineage and predator treatment (presence/absence) leading to a total of 240 replicates. Thirty additional replicates without aphids and ladybeetles were performed as an insect-free control. As it was not possible to perform all replicates simultaneously, we conducted the experiment at three different dates with 6 or 7 of the replicates of each treatment. For each date, we used the same methods and standardization of ladybeetle, aphid and plant age/stage/size.

### Statistical analyses

We performed the statistical analyses in two steps to (1) investigate whether trophic cascade strength differed among aphid lineages to test for the existence of intra-specific differences, and (2) determine whether the observed variation was linked to aphid biotypes to test for the role of host plant specialization. For the first step, we analyzed the effects of predators, aphid lineage, and their interactions on plant fresh aboveground mass and height using Linear Mixed Models (LMMs) with experimental dates added as random effect. A significant and positive predator effect would indicate a significant trophic cascade where plant biomass is higher in presence of predators than in their absence. Moreover, a significant interaction between predator treatment and aphid lineage would indicate that trophic cascade strength (i.e. the effect of predators on plants) would differ among aphid lineage. We next analyzed the effects of predators, aphid lineage, and their interactions on aphid density using a Generalized Linear Mixed Model (GLMM) with a Poisson distribution and a log link function, with experimental dates added as random effect. Finally, we analyzed the effects of aphid lineage on predator fresh body mass using an LMM with experimental dates added as random effect. The significances of the model fixed terms were assessed using Chi-tests from analyses of deviances, and post-hoc Tukey tests were used to determine significant differences among means.

For the second step, we investigated the effects of aphid biotype, predators and their interaction on plant biomass, plant height, and aphid density using LMM and GLMM models as described above but adding lineage identity as a random effect. We also analyzed the effects of aphid biotype on ladybeetle larva body mass using LMM as described above but adding lineage identity as a random effect. Aphid lineage color or its interaction with the two other independent variables did not significantly affect the response variables (P > 0.05) and was thus excluded from final analyses.

To better understand the links between plant response and aphid response to predators, we calculated, for each aphid lineage, the trophic cascade strength defined as the log response ratio of plants to predators: R_p_ = ln(x_P_/x_C_), where x_P_ and x_C_ are the mean values of the plant trait (biomass or height) in the treatment with and without predators, respectively (Hedges, Gurevitch & Curtis 1999). We also calculated this ratio for aphid density (Ra) using the same formula. We next estimated the variance of each log ratio estimate as var(R) = s_P_^2^/(n_P_x_P_^2^)+s_C_^2^/(n_C_x_C_^2^), where n and s respectively denote the number of replicates and the standard deviation in the treatments with predators P and without predators C (Hedges, Gurevitch & Curtis 1999). We then calculated the 95% confidence intervals by multiplying var(R) by 1.96 assuming a normal distribution (Hedges, Gurevitch & Curtis 1999). A log ratio value that does not differ significantly from zero (i.e. when its 95% confidence intervals overlap with zero) indicates the absence of predator effects whereas a positive or negative log ratio value represents a positive or negative effect of predator on the lower trophic level (aphids or plants), respectively (Hedges, Gurevitch & Curtis 1999). A positive log ratio value would thus indicate a trophic cascade where plants benefit from predator presence. It is thus possible to compare the strength of trophic cascades by comparing the log response ratios of plants to predators across treatments.

To evaluate whether the effects of predators on plants and on aphids are positively related, we plotted herbivore density log ratios against plant (biomass or height) log ratios. We next used a linear least squares regression model to investigate the relationship between the direct effects of predators on aphids and their indirect effects on plants. Finally, we investigated the effects of aphid lineage population growth rate on the plant log ratios, aphid log ratios and average predator body mass using linear regression models. The population growth rate of each aphid population (in the absence of predators) was calculated as ln(N_t_/N_0_)/t where N_0_ is the initial aphid density (i.e. 6), N_t_ is the final aphid density and t is the number of experimental days (i.e. 10).

To investigate the relationship between aphid biotype and the strength of predator effects on aphids and plants, we used the raw data from each replicate to calculate for each aphid biotype the mean aphid biotype population growth rate as well as the plant and herbivore log ratios and their variances. We next plotted biotype log ratios against plant log ratios, biotype population growth rate against plant log ratios, biotype population growth rate against biotype log ratio, and biotype population growth rate against predator mass. We considered that log ratios differ significantly if their 95% CIs do not overlap (Hedges, Gurevitch & Curtis 1999). LMMs and GLMM were computed using the lme4 package (Bates et al. 2015), and analyses of deviance were performed using the car package (Fox & Weisberg 2011). In cases interaction terms of the LMMs or GLMM were non-significant, they were removed prior to calculating significance values on the main effects or other interactions. All analyses were performed using R 3.1.1 (R Development Core Team 2017).

## Results

### Influence of aphid lineage and biotype on predator body mass, herbivore density, and plant biomass and height

Ladybeetle body mass depended on aphid lineage (χ^2^ = 55.14, *df* = 5, *P* < 0.0001). Ladybeetle larvae feeding on the aphid lineage LSR1 were about two times heavier than those feeding on the lineage T734 (Fig. 1a). Ladybeetle body mass significantly differed between aphid biotypes, with ladybeetles feeding on aphids of the *Alfalfa* biotype being heavier than those feeding on the *Clover* biotype (χ^2^ = 6.73, *df* = 1, *P* = 0.009, Fig 1b).

**Figure. 1.**
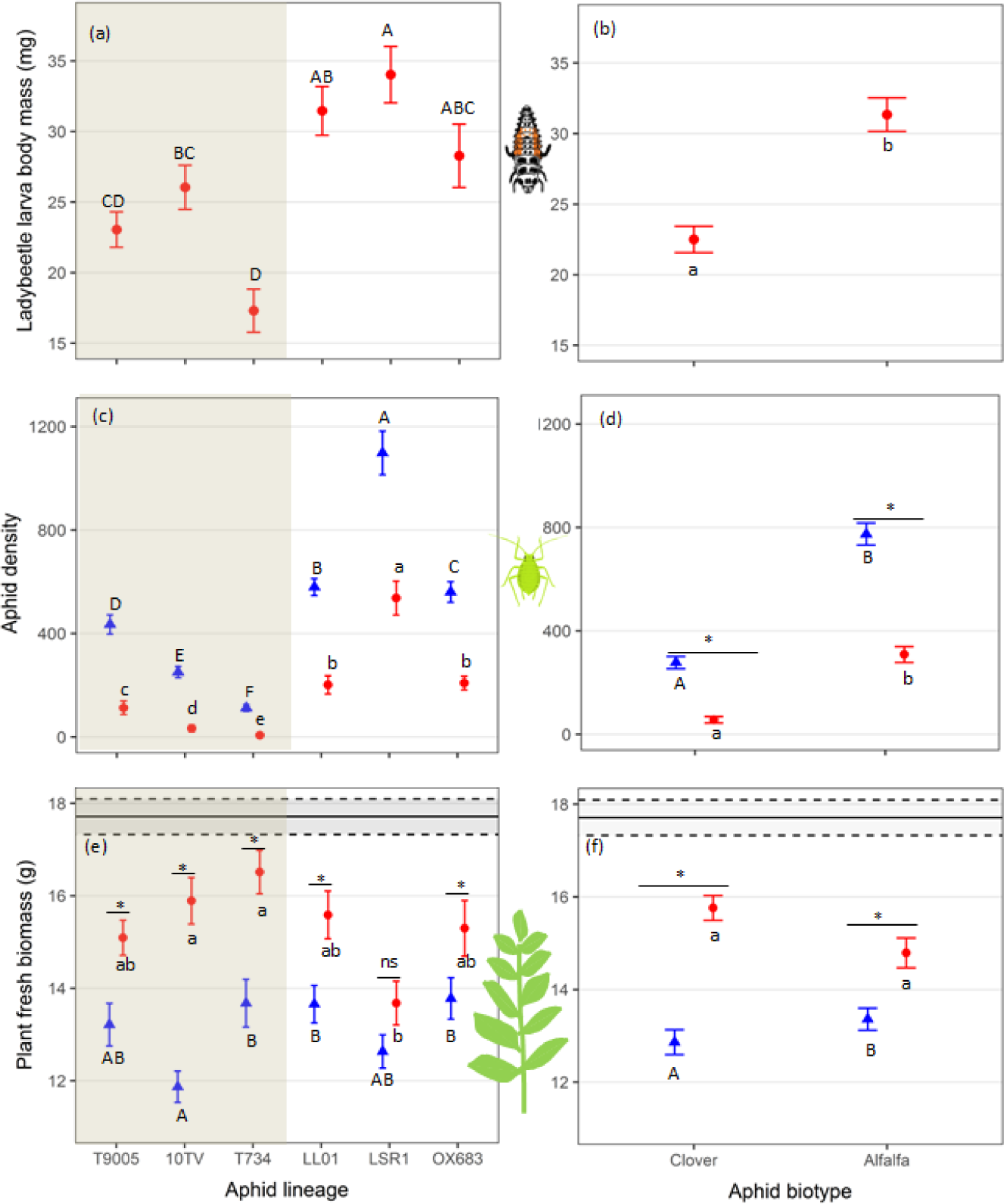
Influence of aphid lineage and biotype on the three trophic levels. <u>Left panels</u>: effects of the six aphid lineages (X axis) on mean (± SE) ladybeetle larva body mass (a), aphid density (c) and plant fresh biomass (e) (*n* = 20 replicates per treatment). Shaded area: aphid lineages of the *Clover* biotype; Non-shaded area: lineages of the *Alfalfa* biotype. <u>Right panels</u>: effects of aphid biotype (X axis) on mean (± SE) ladybeetle larva biomass (b), aphid density (d) and plant fresh biomass (f) (*n* = 60 replicates per treatment). Red dots: with predators; Blue triangles: without predators. Within each panel, small or capital letters denote significant differences (*P* < 0.05) among aphid lineages (panels a, c, e) or between aphid biotypes (panels b, d, f) within each predator treatment. For each lineage or each aphid biotype, an asterisk or “ns” denotes significant (*P* < 0.05) or non-significant (*P* > 0.05) predator effect (significance levels estimated with post hoc Tukey tests), respectively. Black lines of panels e and f represent mean (± SE; dotted lines) plant fresh biomass in controls without aphids or ladybeetles.

Aphid density varied strongly among lineages (χ^2^ = 27370.6, *df* = 5, *P* < 0.0001) with the highest density for the LSR1 lineage and the lowest for the T734 lineage (Fig. 1c). Predators always significantly decreased aphid density (Fig. 1c; χ^2^ = 15175.4, *df* = 1, *P* < 0.0001) although the strength of this effect varied among lineages (significant interaction between lineage and predator treatment: χ^2^ = 1827.9, *df* = 5, *P* < 0.0001).

Aphid density significantly differed between biotypes (χ^2^ = 10.99, *df* = 1, *P* = 0.0009), and was affected by the presence of predators (χ^2^ = 15900.27, *df* = 1, *P* < 0.0001) and by the interaction between predators and biotype (Fig 1d; χ^2^ = 1180.33, *df* = 1, *P* < 0.0001). Aphid density was higher and predator effect on aphid density stronger for the *Alfalfa* biotype than for the *Clover* biotype (Fig. 1d).

Plant biomass significantly varied among aphid lineages (Fig. 1e; χ^2^ = 23.43, *df* = 5, *P* = 0.0003), and was affected by the presence of predators (χ^2^ = 66.72, *df* = 1, *P* < 0.0001) and by the interaction between these two factors (Fig 1e; χ^2^ = 13.57, *df* = 5, *P* = 0.0185). Without predators, lineage 10TV had a stronger impact on plant fresh biomass than T734, LL01 or OX683, whereas with predators, lineages 10TV and T734 had a weaker impact on plant fresh biomass than LSR1 (Fig. 1e). The significant interaction between predator treatment and aphid lineage indicates that the effect of predators on plant biomass (i.e. trophic cascade strength) depended on aphid lineage. Post-hoc tests indicated that predators indirectly increased plant biomass but this increase depended upon the lineage with a large effect for 10TV and T734 and a weak non-significant one for LSR1 (Fig. 1e and Table S1).

Plant biomass was significantly influenced by the predator treatments (Fig. 1f; χ^2^ = 65.58, *df* = 1, *P* < 0.0001), and by the interaction between predator treatments and aphid biotypes (χ^2^ = 6.8851, *df* = 1, *P* = 0.0087). Without predators, the *Clover* biotype had a stronger impact on plant biomass compared to the *Alfalfa* biotype (Fig. 1f, blue dots). The positive indirect effect of predators on plant biomass was stronger in plants exposed to the *Clover* than to the *Alfalfa* aphid biotypes (Fig. 1f, differences between red and blue dots, Table S2). The effects of aphid lineage or biotype, predators and their interactions on plant height were qualitatively similar than their effects on plant biomass (see Fig. S1 and Text S1 for more details).

### Relationship between the effects of predators on plants and their effects on aphids

Predator direct effect on aphid density (i.e. herbivore density log ratio, *X* axis in Fig. 2) was always significant as indicated by the non-overlap of log ratio confidence intervals with the intercept (plain vertical black line in Fig. 2a). The magnitude of this predator effect differed among aphid lineages and was minimal for the LSR1 lineage and maximal for the T734 lineage. Interestingly, aphid biotypes influenced the predator direct effects on aphid density, which was stronger for the *Clover* than the *Alfalfa* biotype (Fig. 2b).

**Figure. 2.**
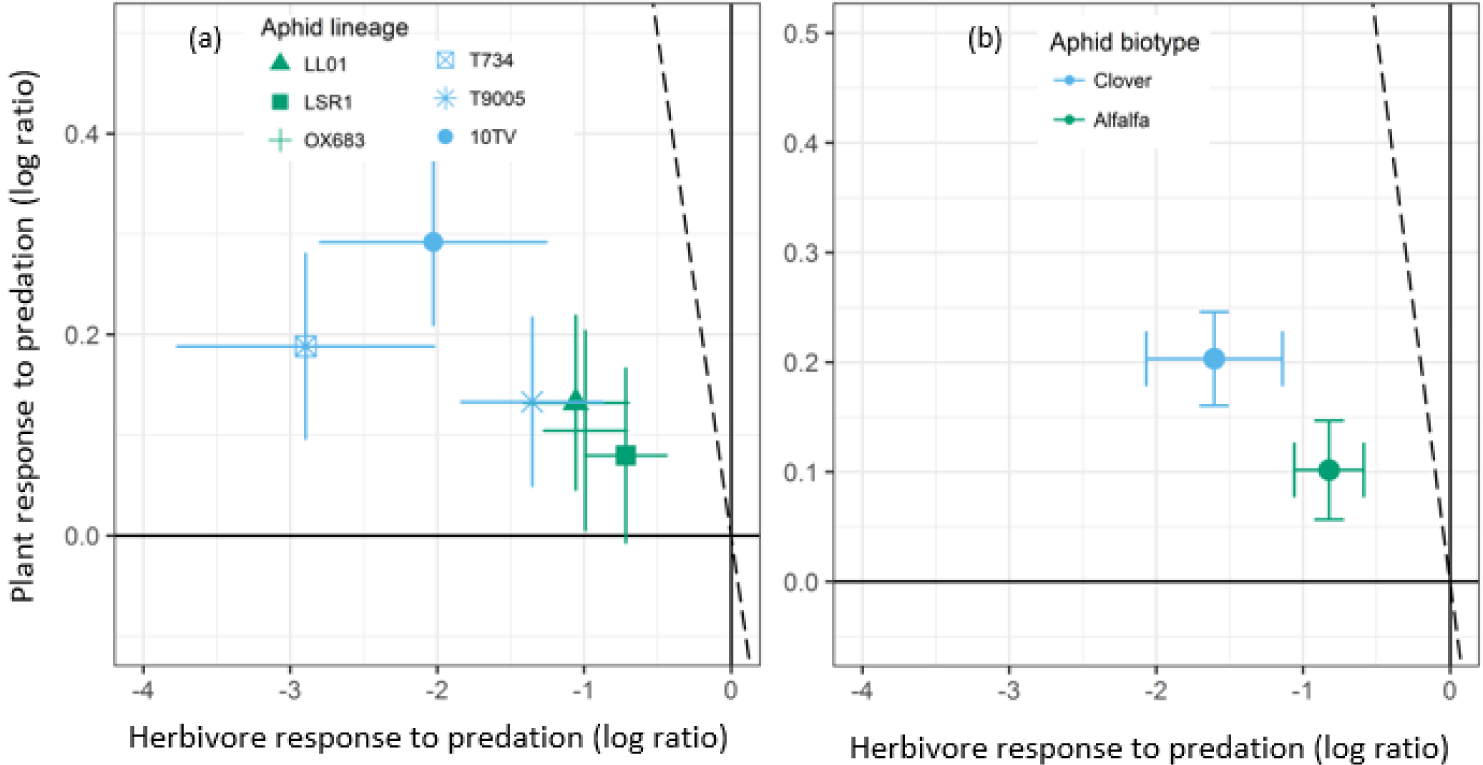
The relationship between the magnitude (log ratio ± 95% CI) of the direct effect of predator on herbivore density (i.e. herbivore response to predation) and the indirect effect of predator on fresh biomass of plants (i.e. plant response to predation) by aphid lineage (a) and biotype (b). Predator effect is significant if the confidence interval does not overlap zero (dark full lines). For plant response to predation, a significant trophic cascade corresponds to values above the horizontal line. For herbivore response to predator, a significant impact of predators on aphid population corresponds to values on the left side of the vertical line. The dotted line shows the 1:1 relationship, representing equivalence of predator direct and indirect effects. If the data cluster to the left of the 1:1 line, then top-down effects are attenuating at the plant level; if they cluster to the right of the 1:1 line, then top-down effects are intensifying and, if they cluster along the 1:1 line, the effect magnitudes do not attenuate.

The indirect effect of predators on plant biomass varied significantly among lineages and differed from zero except for the LSR1 lineage (Fig. 1a and Fig. 2a). Moreover, predator indirect effect on plant biomass was significantly stronger for the *Clover* than for the *Alfalfa* biotype (Fig. 2b). The relationship between predator effects on plant biomass and on aphids was non-significant (F_(1,4)_ = 3.80, *P* = 0.12, *R*^*2*^ = 0.36 Fig 2a). Finally, all data point cluster to the left of the 1:1 dotted line indicating strong attenuation of top-down effects at the plant level. The predator indirect effect on plant height as well as the influence of aphid lineages and biotype on this effect were qualitatively similar to these obtained for the plant biomass (see Fig. S2 and Text S2 for more details).

### Relationship between aphid population growth rate and predator effects on plants

Although predator indirect effects on plant biomass tended to decrease with lineage population growth rate (Fig 3a), this relationship was non-significant (Fig. 3; F_(1,4)_ = 3.68, *P* = 0.12, *R*^*2*^ = 0.35). Interestingly, predator indirect effects on plant biomass were stronger with *Clover* than with *Alfalfa* biotype despite the faster population growth rate of the later (Fig. 3b). The results for plant height were qualitatively similar to those obtained for plant biomass (see Fig. S2 and Text S2 for more details).

**Figure. 3.**
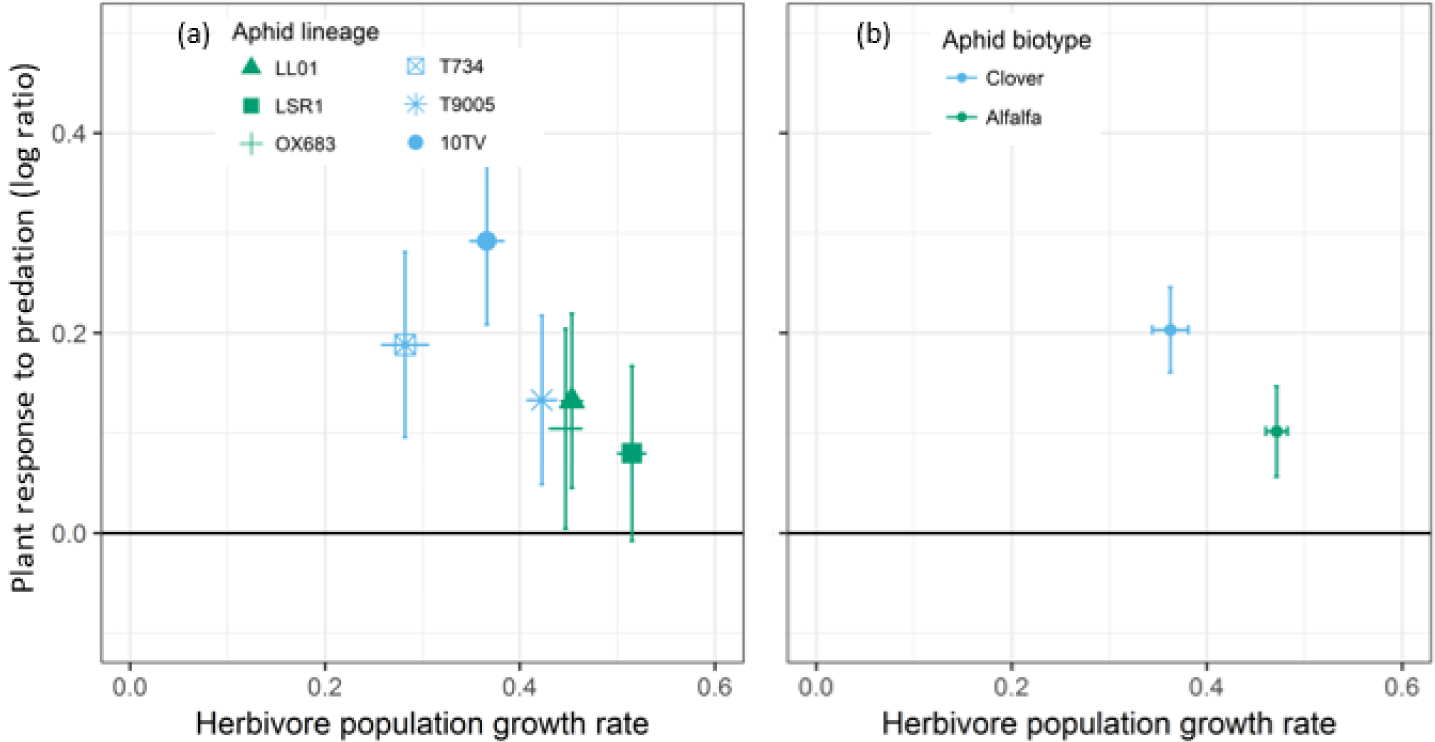
Relationship between aphid population growth rate (mean ± 95% CI) and the magnitude (log ratio ± 95% CI) of predator indirect effect on plant fresh biomass according to aphid lineage (a) and biotype (b). Predator effect is significant if the 95% CI does not overlap the X axis (dark full line).

### Influence of aphid population growth rate on herbivore density log ratio and on predator body mass

The effect of predators on aphid density (i.e. herbivore log ratio) was associated with aphid lineage population growth rate (F_(1,4)_ = 132.96, *P* = 0.000323, *R*^*2*^ = 0.96; *y* = 9.94*x-*6.62, Fig. 4a) showing that predators have a weaker effect on fast growing aphid lineages than on slow growing aphid lineages. Predator body mass was positively associated with aphid lineage population growth rate (F_(1,4)_ = 20.29, *P* = 0.01079, *R*^*2*^ = 0.79; *y* = 68.14 *x-*1.54, Fig. 4c) indicating that lineages with fastp opulation growth result in larger predators than lineages with slow population growth. Interestingly, lineages of the *Clover* and *Alfalfa* biotypes clustered separately along the regression lines in figure 4a and 4c indicating that the influence of biotype on predator effect on aphids and on predator body mass are mainly linked to differences between biotype population growth rates. Grouping the data by aphid biotype (Fig. 4b and d) confirmed that population growth rate, predator effect on aphid density, and predator body mass differed between the two biotypes.

**Figure. 4.**
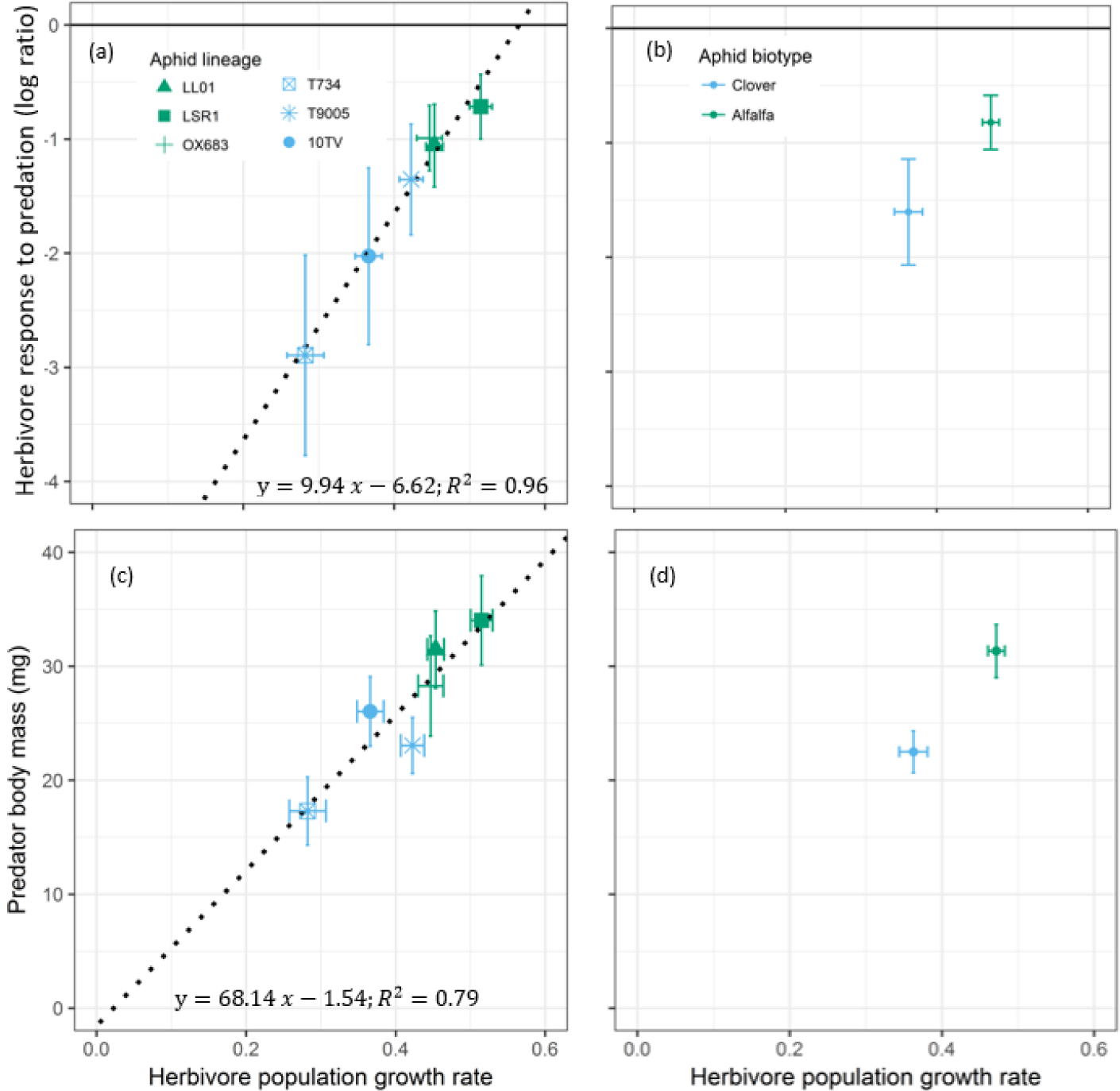
Relationship between aphid population growth rate in the absence of predators (mean ± 95% CI) and (first row) the direct effect of predators on aphid density (log ratio ± 95% CI) and (second row) predator body mass (mean ± 95% CI) according to aphid lineage (a, c) and biotype (b, d). In panel a and b, predator effect is significant if 95% CIs do not overlap the X axis (dark full line).

## Discussion

Although intraspecific differences in organism phenotype and behaviour have clear implications for pairwise species interactions, their effects on higher order interactions remain largely unexplored (Bolnick et al. 2011; Toscano & Griffen 2014; Belgrad & Griffen 2016; Sanders et al. 2016). Here, we quantified the impact of pea aphid clonal lineages (specialized on alfalfa or clover) on their universal legume host plant Vicia fabae in the presence or absence of a ladybeetle predator. We showed that the strength of trophic cascade strongly depends on intraspecific differences among herbivores and their host-plant specialization. Our study thus highlights the importance of intraspecific differences and host-plant specialization as drivers of the strength of trophic cascades.

### Effects of herbivore lineages on trophic cascade strength and predator body mass

We found that on average, predators decreased aphid population density by 66.03 % (± 0.24 %, 95 % CI), which, in turn, increased plant biomass by 16.29 % (± 3.72 %) and plant height by 20.18 % (±3.07 %). These values are within the range of values reported by previous meta-analyses on trophic cascades in terrestrial system (Schmitz et al. 2000; Shurin et al. 2002; Borer et al. 2005; Romero & Koricheva 2011) and confirm the previously described strong attenuation of the predator top-down effect down the food chain (Polis & Strong 1996; Borer et al. 2005). Nevertheless, our study indicates that considering only these average values limits our understanding of multi-trophic interactions as we found that the strength of trophic cascade strongly depends on herbivore lineages with cascading effect of predator on plants ranging from non-significant (0%) to a significant increase of 34% in plant biomass (Figs. 1 and 2). These strong differences in the strength of trophic cascade mediated by herbivore intraspecific trait variation could contribute to explain why (1) previous studies had difficulties in assessing the strength and occurrence of trophic cascades (Schmitz et al. 2000; Halaj & Wise 2001), (2) the occurrence and strength of trophic cascades strongly differ among studies, species, and habitats (Schmitz et al. 2000; Bell, Neill & Schluter 2003; Borer et al. 2005), and (3) no single hypothesis can explain variation in the magnitude of trophic cascades (Borer et al. 2005).

Interestingly, intraspecific differences among herbivore lineages did not only influence top-down effects but also climbed up the food chain and influenced the predator phenotype. Indeed, we found that predator body mass depends on which aphid clonal lineage and biotype they are feeding on (Fig. 1). To the best of our knowledge, this is the first experimental evidence of a predator body mass being significantly influenced by the intraspecific specialization of herbivore prey on particular host-plants. Our results indicate that this effect is likely driven by the aphid population growth rate that strongly differs between biotypes: predators were larger on fast growing aphid lineages (Fig. 4). Nevertheless, different traits such as inter-biotype variation in defensive behavior or palatability may also contribute to explaining the effects of aphid biotype on predator body mass (Kunert et al. 2010; Ben-Ari et al. 2019). Body size is a key trait that determines many ecological properties including fecundity, behaviour, population growth rate, trophic position, species interactions and community stability (Peters 1983; Brose et al. 2006; White et al. 2007). This implies that the effects of herbivore intraspecific variation and ecological specialization on predator body mass are likely to influence predator populations and thereby have long-term effects on the dynamics and structure of the community.

### Investigating the mechanisms underpinning the influence of intraspecific variation on trophic cascade

An important step toward a better understanding of trophic cascade functioning is to explain how intraspecific differences at a given trophic level can influence adjacent trophic levels as well as predator indirect effects on plants (i.e. trophic cascade strength). As the conventional view is that the strength of trophic cascade strongly depends on the density of the interacting species (Schmitz et al. 2000; Schmitz, Krivan & Ovadia 2004; Borer et al. 2005), we hypothesized that differences in the population growth rate of aphid lineages would explain the intensity of lineages’ impact on plant, predators, and the strength of trophic cascades. Accordingly, the predator direct effect on aphid density strongly depended on the lineages’ population growth rate with fast growing lineages being less impacted by predators than slow growing lineages (Fig. 4). Differential population growth rate among aphid biotypes thus explains why (1) the direct effect of predators on herbivore density is weaker for the Alfalfa than for the Clover biotype, and (2) ladybeetle larvae reach a larger body mass when feeding on the Alfalfa than on the Clover biotype (as mentioned above). We thus conclude that the ladybeetle-aphid interaction is strongly density-dependent and that the differential effects of aphid lineages or biotypes on this interaction are mainly linked to their differential population growth rate. The observation that alfalfa lineages reach higher densities on the broad bean (in absence of predators) suggests that the alfalfa biotype performs better on the broad bean than the clover biotype, in agreement with a previous study (McLean et al. 2010). It would be interesting to conduct a similar experiment on the three host plants (clover, alfalfa and broad bean) to test this hypothesis and investigate potential trade-offs in host-plant specialization and their consequences for predator traits and the strength of trophic cascades.

On the other hand, the direct effect of aphid lineages on plant biomass and height was not related to their population growth rate but was instead mainly linked to their host plant specialization. Surprisingly, plants were more impacted by the Clover than the Alfalfa biotype despite the faster population growth of the latter (Fig. 3). This counter-intuitive result contradicts the herbivore efficiency hypothesis predicting that herbivores with the highest population growth rate should have the strongest effect on plants, which in turn should increase trophic cascade strength when predators are efficient in reducing herbivore populations (Borer et al. 2005). The differential effects of aphid biotype on plants could be linked to morphological, physiological and behavioural differences between aphid biotypes (Via 1991; Kunert et al. 2010) and/or linked to the plant defensive response against a given aphid biotype (Via 1991; Tétard-Jones et al. 2007). For instance, biotype-specific aphid effectors injected while feeding may be recognized differentially by the host plants and trigger more or less defense responses (Boulain et al. 2019). It is also possible that Clover biotypes feed and impacts more the plant than the Alfalfa biotypes but has a lower assimilation efficiency leading to a reduced population growth despite a strong effect on the plant.

Whatever the exact mechanism driving the differential impact of aphid biotype on plants, we found that the strength of trophic cascade strongly depends on herbivore biotypes and lineages and is not directly related to the predator effect on aphid density. Indeed, we found no straightforward relationship between the direct effect of predators on herbivore density and their indirect effect on plant traits. This indicates that herbivore intraspecific differences and host-plant specialization play a stronger role in determining trophic cascade strength than the density-dependent effects related to herbivore population growth rate. More generally, herbivore intraspecific differences induced considerable changes in our tritrophic system that could not be predicted from observations on a bitrophic system. We thus conclude that going beyond pairwise interactions and considering the links between intraspecific trait variation and evolutionary divergence associated to host-plant specialization is certainly a promising avenue to better understand multitrophic interactions.

### Ecological and evolutionary implications of herbivore intraspecific trait variation

Herbivores link primary producers with higher trophic levels. Variation in herbivore traits can thus have important consequences for the dynamics of ecological communities as shown by previous studies focusing on pairwise interactions (Bolnick et al. 2011; Sentis, Morisson & Boukal 2015). Our results indicate that intraspecific variation in herbivore lineages and their ecological specialization can also have important consequences for higher-order interactions and trophic cascade strength. While the duration of our study was too short to measure feedback loops, we expect that the contrasting effects of aphid biotype on plants and predators may feedback and have a long-lasting effects on predator and prey populations. For instance, smaller ladybeetles lay fewer (Dixon & Guo 1993; Bista 2013) and smaller eggs (Osawa 2005; Kajita & Evans 2010), which should in turn reduce top-down pressure, thereby allowing for the larger growth of aphid populations. As a result, we would then expect a stronger impact on plants which would then feedback on herbivore populations. We thus argue that herbivore trait variation is likely to affect population dynamics on the longer term and should thus receive more attention to better understand the structure and dynamics of ecological communities. More generally, intraspecific variation at any trophic level might be influential and further studies are needed to determine when and where intraspecific variation has the strongest influence on trophic cascades.

### Conclusion

Intraspecific variation is central to our understanding of evolution and population ecology, yet its consequences for community ecology are poorly delineated (Bolnick et al. 2011; Violle et al. 2012). Here, we showed that intraspecific differences among herbivore lineages influences the strength of trophic cascades. Interestingly, differences in the strength of trophic cascades were more related to aphid lineage and host-plant specialization than to density-dependent effects mediated by the growth rate of aphid populations. Our findings imply that intraspecific trait diversity and host-plant specialization are key drivers of the strength of trophic cascades and therefore they should not be overlooked to decipher the joint influence of evolutionary and ecological factors on the functioning of multitrophic interactions.

## Data accessibility

Data are available online: DOI: 10.5281/zenodo.3716708

## Supplementary material

Script and codes are available online: DOI: 10.5281/zenodo.3718961

## Acknowledgements

We thank Sara Magalhães, Bastien Castagneyrol, and an anonymous reviewer for their positive and constructive comments. This work was supported by ANR funded French Laboratory of Excellence projects ‘LABEX TULIP’ and ‘LABEX CEBA’ (ANR-10- LABX-41, ANR-10-LABX-25-01) and ANR funded Toulouse Initiative of Excellence “IDEX UNITI” (ANR11-IDEX-0002-02). AS was also founded by the People Program (Marie Curie Actions) of the European Union’s Seventh Framework Program (FP7/2007-2013) under REA grant agreement n°PCOFUND-GA-2013-609102, through the PRESTIGE program coordinated by Campus France. Version 4 of this preprint has been peer-reviewed and recommended by Peer Community In Ecology (https://doi.org/10.24072/pci.ecology.100047)

## Author contributions

A. S., R. B., E. D. and J-L. H. conceived and designed the experiments. A.S., R. B., N. D., and A. M. performed the experiments. A. S. analyzed the data and wrote the first draft of the manuscript. A. S., J-C. S., A. M., B. P., E. D., and J-L. H. contributed substantially to revisions.

## Appendix

**Table S1.**
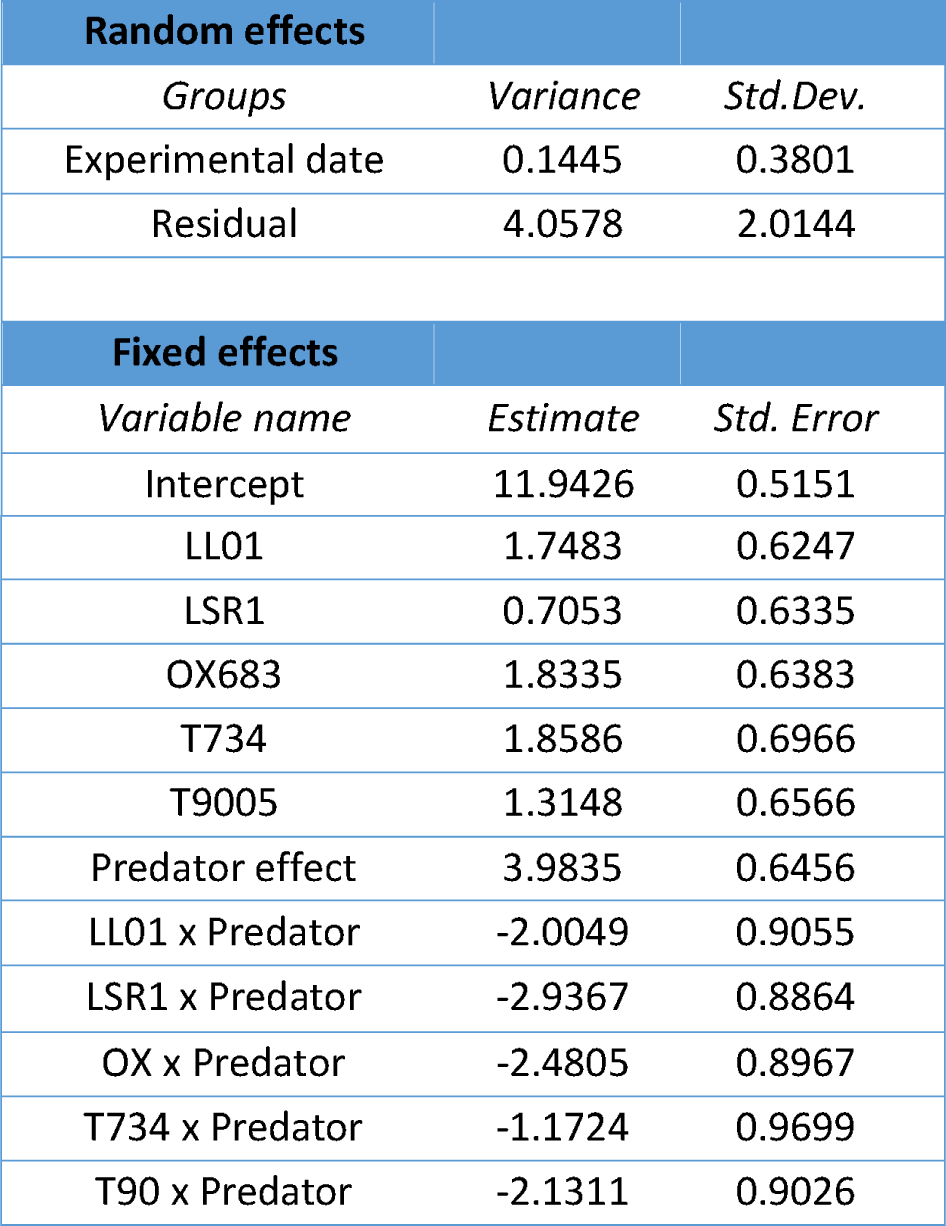
Values of the variance and standard deviation for the random effects and values of the estimate, and standard error for the fixed effects of the LMMs for the effects of aphid lineage, predator presence, and their interactions on plant biomass.

| Random effects |  |  |
| --- | --- | --- |
| *Groups* | *Variance* | *Std.Dev.* |
| Experimental date | 0.1445 | 0.3801 |
| Residual | 4.0578 | 2.0144 |
| Fixed effects |  |  |
| *Variable name* | *Estimate* | *Std. Error* |
| Intercept | 11.9426 | 0.5151 |
| LL01 | 1.7483 | 0.6247 |
| LSR1 | 0.7053 | 0.6335 |
| OX683 | 1.8335 | 0.6383 |
| T734 | 1.8586 | 0.6966 |
| T9005 | 1.3148 | 0.6566 |
| Predator effect | 3.9835 | 0.6456 |
| LL01 x Predator | -2.0049 | 0.9055 |
| LSR1 x Predator | -2.9367 | 0.8864 |
| OX x Predator | -2.4805 | 0.8967 |
| T734 x Predator | -1.1724 | 0.9699 |
| T90 x Predator | -2.1311 | 0.9026 |

**Table S2.** Values of the variance and standard deviation for the random effects and values of the estimate, and standard error for the fixed effects of the LMMs for the effects of aphid biotype, predator presence, and their interactions on plant biomass.

| Random effects |  |  |
| --- | --- | --- |
| *Groups* | *Variance* | *Std. Dev.* |
| aphid lineage | 0.4586 | 0.6772 |
| date | 0.1348 | 0.3671 |
| Residual | 4.1077 | 2.0267 |
| Fixed effects |  |  |
| *Variable name* | *Estimate* | *Std. Error* |
| Intercept | 12.9829 | 0.5265 |
| Alfalfa biotype | 0.3863 | 0.6678 |
| Predator effect | 2.8710 | 0.3854 |
| Alfalfa x Predator | -1.3855 | 0.5280 |

### Text S1. Influence of aphid lineage and biotype on plant height

Plant height varied among aphid lineages (χ^2^ = 80.45, *df* = 5, *P* < 0.0001), predator treatments (χ^2^ = 49.94, *df* = 1, *P* < 0.0001), and the interaction between aphid lineage and predator treatments (χ^2^ = 20.04, *df* = 5, *P* = 0.0012). Without predators, lineages 10TV and LSR1 showed the strongest impact on plant height whereas, with predators, lineage T734 had the weakest impact (Fig. S1a). Predator presence increased plant height but the magnitude of this effect varied significantly among aphid lineages (Fig. S1a).

Plant height was significantly influenced by the interaction between predator treatments and biotypes (χ^2^ = 13.71, *df* = 1, *P* = 0.0002). Without predators, plant height diminished more for the *Clover* than the *Alfalfa* biotypes (Fig. S1b). With predators, plant height did not significantly differ between biotypes (Fig. S1b). Finally, the predator positive effect on plant height was stronger for plant exposed to the *Clover* than to the *Alfalfa* biotypes (Fig. S1b).

**Figure S1.**
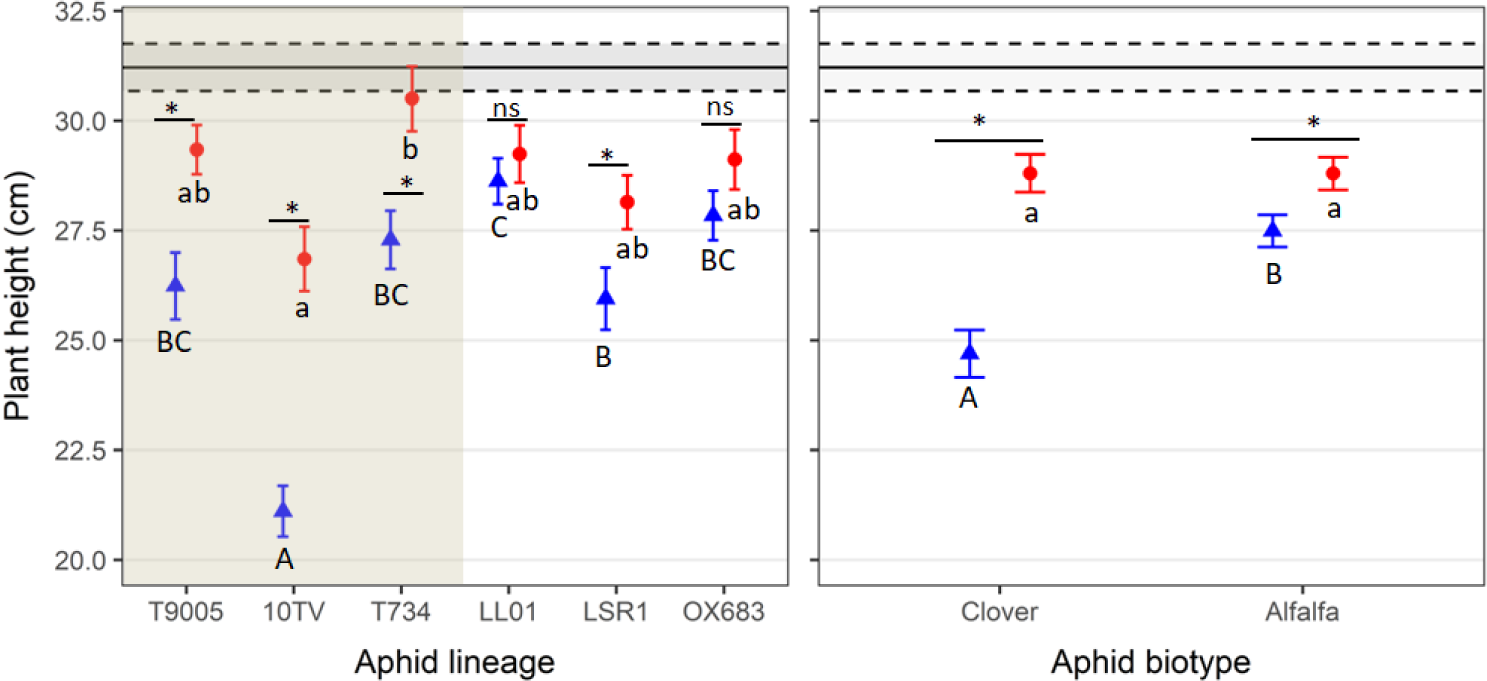
Plant height (mean ± SE) with (red dots) or without (blue triangles) predators according to aphid lineage (a) and biotype (b). Within each panel, letters denote significant differences (*P* < 0.05) among aphid lineages (panel a) or aphid biotypes (panel b) with predators (small letters) or without predators (capital letters). Asterisk or “ns” denote significant (*P* < 0.05) or non-significant (*P* > 0.05) effect of predators within each lineage in (a) or aphid biotype in (b). Black lines represent mean (± SE; dotted lines) plant height in controls without aphids or ladybeetles.

### Text S2. Relationship between the effects of predators on plant height and their effects on aphids

Predator indirect effect on plant height varied significantly among lineages and was not always significantly different from zero as for the OX683 and LL01 lineages (Fig. S2a). Moreover, predator indirect effect on plant was significantly stronger for the *Clover* than for the *Alfalfa* biotype (Fig. S2b). The relationship between predator effects on plants and on aphids was non-significant (F_(1,4)_ = 1.55, *P* = 0.28, *R*^*2*^ = 0.10). Finally, all data point cluster to the left of the 1:1 dotted line indicating strong attenuation of top-down effects at the plant level.

**Fig. S2.**
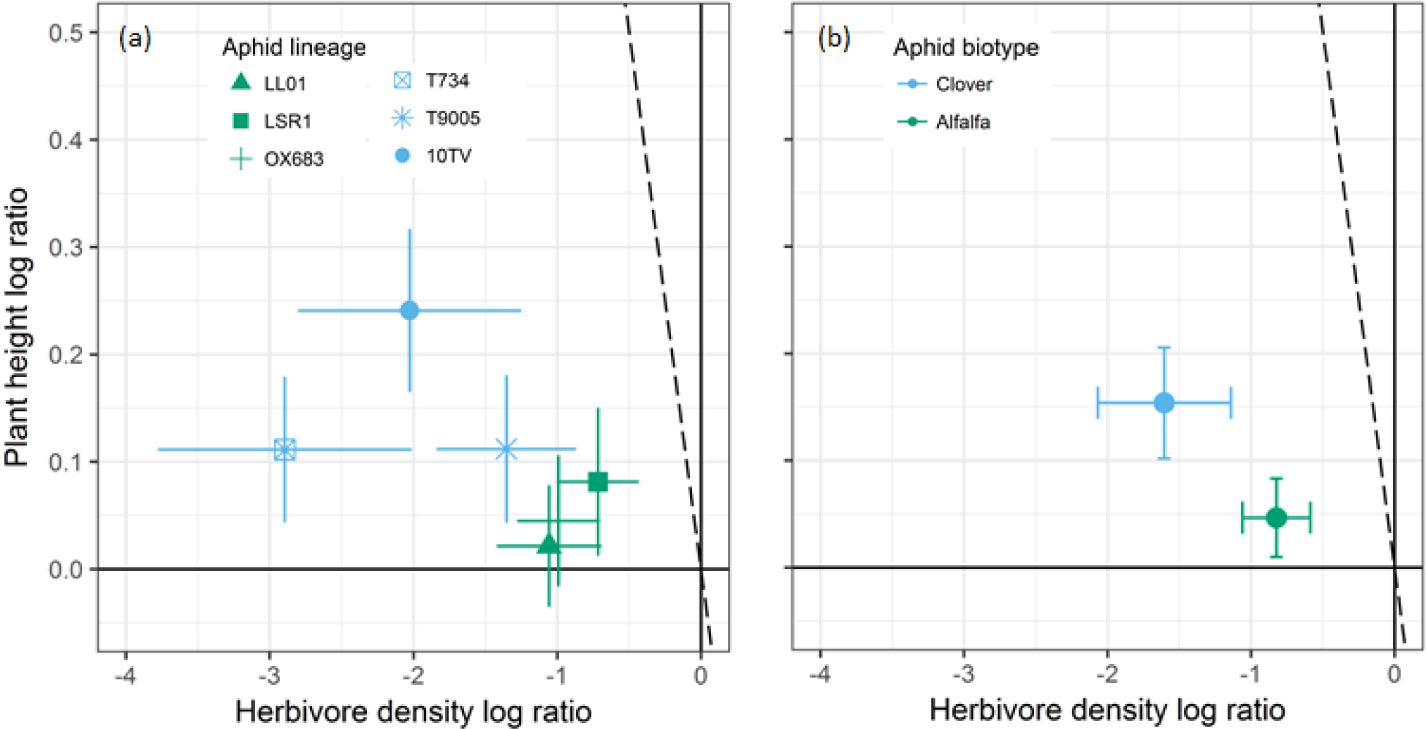
Relationship between the magnitude (log ratio ± 95% CI) of the predator effects on aphid density and on plant height according to aphid lineage (a) and biotype (b). Predator effect is significant if the confidence interval does not overlap zero (dark full lines). The dotted line shows the 1:1 relationship, representing equivalence of predator direct and indirect effects. If the data cluster to the left of the 1:1 line, then top-down effects are attenuating at the plant level; if they cluster to the right of the 1:1 line, then top-down effects are intensifying and, if they cluster along the 1:1 line, the effect magnitudes do not attenuate.

### Text S3. Relationship between aphid population growth rate and predator effects on plant height

Although predator indirect effects on plant height tended to decrease with lineage population growth rate (Fig S3a), this relationship was non-significant (F_(1,4)_ = 1.20, *P* = 0.33, *R*^*2*^ = 0.04). Interestingly, predator indirect effects on plant height were stronger with *Clover* than with *Alfalfa* biotype despite the faster population growth rate of the later (Fig. S3b).

**Fig. S3.**
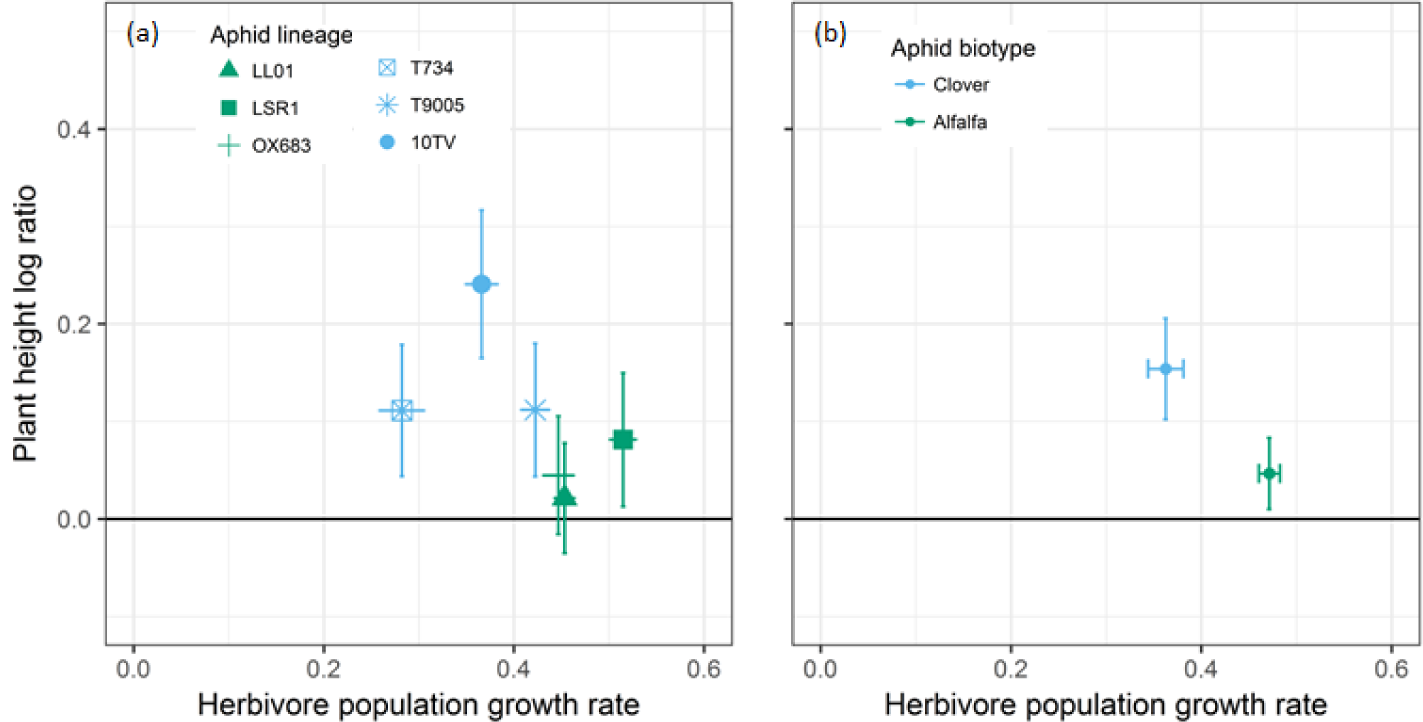
Relationship between aphid population growth rate (mean ± 95% CI) and the magnitude (log ratio ± 95% CI) of predator indirect effect on plant height according to aphid lineage (a) and biotype (b). Predator effect is significant if the 95% CI does not overlap the X axis (dark full line).

